# Transverse Sheet Illumination Microscopy

**DOI:** 10.1101/2025.03.14.642160

**Authors:** Javier Carmona, Blake Madruga, Steve Mendoza, Brian Jeong, Laurent Bentolila, Katsushi Arisaka

## Abstract

Recording the neural activity of biological organisms is paramount in understanding how they process the world around them. Fluorescence microscopy has served as the standard in recording this neural activity due to its ability to capture large populations of neurons simultaneously. Recent efforts in fluorescence microscopy have been concentrated in imaging large scale volumes, however, most of these efforts have been limited by spatiotemporal and bandwidth constraints. We present a novel system called Transverse-Sheet Illumination Microscopy (TranSIM), which captures spatially separated planes onto multiple two-dimensional sCMOS sensors at near diffraction limited resolution with 1.0 µm, 1.4 µm, and 4.3 µm (x, y, and z respectively). The parallel use of sensors reduces the bandwidth bottlenecks typically found in other systems. TranSIM allows for the capturing of data at large-scale volumetric field-of-views up to 748 × 278 × 100 µm^3^ at 100 Hz. Moreover, we were able to capture smaller field-of-views of 374 × 278 × 100 µm^3^ at a faster volumetric rate of 200 Hz. Additionally, we found that the system’s versatile design allowed us the ability to change the vertical magnification programmatically rather than necessitating a change of objectives. With this system we were able to observe intricate communication between neuron populations separated by vast three-dimensional distances, raising the potential to answer complex questions in Neurobiology.

## Introduction

To better understand how animals process the world, neuroscientists observe and analyze their neuronal activity. Through the lens of microscopy, researchers have been able to capture this neuronal activity with unparalleled spatial resolution since the introduction of genetically encoded calcium indicators (GECIs)^1–5^. GECIs are now commonplace throughout the field of neuroscience due to their ability transcribed into model organisms and expressed pan-neuronally or selectively down to single neurons^4,6–9^However, capturing complex neuronal activity is still constrained by spatiotemporal limitations that microscopy research. In two dimensions, microscopy inherently performs well, allowing routine speeds to accurately trace neuronal activity in individual planes. Yet, extending this problem to three dimensions often forces compromises on either spatial or temporal information, and sometimes both^10,11^.

These limitations stem from two primary factors: the volumetric field of view, and more critically, the total system bandwidth. The first limitation primarily arises from the need to take multiple narrow depth-of-field (DOF) images produced by microscopes and create composites of these for three-dimensional reconstructions, thereby slashing the volumetric imaging rate by the total number of axial frames desired for the reconstruction. The second of these limitations, and the most relevant, is the total system bandwidth. Cameras have a finite pixel rate, and here lies the true problem in three-dimensional microscopy, which is regardless of whether we scan or not for three-dimensional imaging, we will always be bandwidth limited. For instance, even capturing the brain of Drosophila.melanogaster (fruit fly), spanning 590 × 340 × 120 µm^3^ and housing approximately 100,000 neurons^12–15^ remains challenging with current technology. Therefore, developing tools that enable volumetric imaging with preserved spatial coverage and temporal resolution, without sacrificing either, remains a pressing goal.

To address these challenges, recent advancements in single-photon (1P) and two-photon (2P) systems have offered innovative solutions tailored to overcoming spatiotemporal limitations in volumetric imaging. In 1P systems, oblique imaging techniques have been developed to reduce the need for axial scanning, a key bottleneck in conventional microscopy^16–19^. By tilting sheets of light at an oblique angle, fluorescence can be simultaneously illuminated and detected using the same objective lens, like traditional orthogonal geometry systems. This approach eliminates the need for mechanical movement of the objective, significantly increasing volumetric imaging speeds. State-of-the-art oblique illumination systems have achieved imaging rates exceeding 100 volumes per second, with fields of view around 200 × 257 × 128 µm^3^, leveraging ultra-high-speed cameras with imaging bandwidths reaching approximately 1.2 GPixels/second^19^. However, a notable limitation of such systems is the time required for each illuminated plane to dwell over the sample, which is necessary to minimize motion blur. This trade-off often imposes a constraint on the overall volumetric acquisition rate.

Conversely, 2P systems have been gaining a lot of traction in the field of microscopy due to their ability to image with localized excitation and deep sample penetration^20,21^. They do so by packing ∼100 femtosecond light pulses into a small enough volume to induce non-linear excitation of the fluorescent indicators. This method of excitation inherently makes 2P systems a temporal domain imaging device, requiring each point in a three-dimensional volume to be imaged one at a time. Recent innovations in 2P systems have been aimed at addressing volumetric imaging challenges. Temporal multiplexing techniques have emerged as a promising strategy for addressing the imaging bottleneck^22–25^. By introducing optical delay lines into the illumination path, individual femtosecond laser pulses can be temporally and spatially multiplexed across axial planes. This allows for simultaneous excitation at multiple depths, achieving voxel rates up to 141 MHz^25^. Despite these advances, the detection side of 2P systems faces bandwidth limitations due to reliance on single photomultipliers. These devices are inherently constrained by the fluorescence lifetime of commonly used fluorophores, such as GCaMP (∼6-7 ns for calcium imaging), and the laser repetition rate^25,26^. These factors collectively limit the effective voxel throughput of the system, highlighting the need for alternative detection strategies that can handle parallelized data acquisition.

Light Field Microscopy (LFM) represents another promising approach for high-speed volumetric imaging, offering distinct advantages and trade-offs^27–29^ captures a snapshot of a volumetric field by simultaneously capturing position and angular information of the light field, enabling fast volumetric imaging speeds^30^. This makes LFM well-suited for studying dynamic biological processes such as neuronal activity or cardiac motion. However, LFM’s reliance on computational reconstruction introduces a significant limitation: long reconstruction times. Depending on the system configuration and computational resources, these reconstruction times can range several seconds to a several minutes per volume^27,29,31–33^. Moreover, light-field deconvolution requirement hinders its applicability for real-time imaging scenarios. Nevertheless, there are reported systems reaching an impressive imaging volume with a diameter of 800 μm and a depth range of 180 μm at 400 Hz^34^. However, these speeds come at the substantial cost to spatial resolution, with reported resolutions of 4 × 4 × 12 μm^3^ near the center of the volume to 8 × 8 × 30 μm^3^ on the edges of the volume. Despite these constraints, the simplicity of its optical setup and the potential for parallelized processing make LFM an appealing solution for specific applications where temporal resolution can be balanced against computational overhead.

Lately, there has been innovation in microscopy research showing that parallel detection is possible^35–37^. These systems rely on the placement of deflection mirrors or pinholes along the detection pathway. This coaxial placement of optical components leads to high pass filtering effect akin to a Fourier domain DC component removal. Here, we demonstrate the use of a system that combines the advantages of parallel data acquisition and axial and lateral separation of excitation planes. This system, we named Transverse Sheet Illumination Microscopy (TranSIM), employs a robust yet simple fully reflective mirror and partially reflective mirror combination to generate axially and laterally separated 1P excitation beams. This laser-shearing mechanism allows for spatial demixing of detection light paths, enabling near-diffraction-limited spatial resolution and near-millisecond temporal resolution. TranSIM adopts the same scanning geometry as a line-scanning confocal microscope, where a thin line of focused laser light is swept across the sample to sequentially excite fluorophores within the imaging field^38–42^. Moreover, unlike conventional line-scanning confocal systems that rely on single-plane illumination and detection, TranSIM achieves parallelized illumination and detection across multiple depths. Specifically, the system uses multiple axially and laterally displaced excitation lines that are simultaneously projected into the sample. On the detection side, each illuminated line is imaged in parallel, enabling the system to capture data from all illuminated planes simultaneously. This parallelized approach significantly improves volumetric acquisition speed while maintaining the high spatial resolution associated with line-scanning systems.

By incorporating parallelized depth illumination and detection, TranSIM overcomes the inherent trade-offs faced by traditional line-scanning systems, where achieving higher temporal resolution typically requires sacrificing spatial coverage or resolution. Furthermore, the system eliminates the need for computational reconstruction typical of methods like LFM, as the volumetric data is captured directly in the spatial domain, thus viewable in real-time. Lastly, by cleverly displacing the illumination beams, we can avoid any optical impedance that would otherwise affect the image quality, for example by high pass filtering the images. These capabilities make TranSIM a robust tool for dynamic imaging of large-scale neuronal activity, offering a unique combination of high speed, wide field of view, and near-diffraction-limited resolution.

## Results

The illumination and detection of simultaneous imaging planes are the cornerstone of TranSIM’s design and functionality. By parallelizing the acquisition of imaging planes, TranSIM enables the rapid and efficient capture of volumetric data, significantly enhancing temporal resolution. The system achieves this by creating and detecting multiple imaging planes that are separated both laterally and axially, allowing for a comprehensive 3D representation of the sample in real-time.

This approach eliminates the need for traditional axial scanning mechanisms, which are commonly employed in conventional microscopy systems such as light-sheet and confocal microscopy. In those systems, volumetric imaging requires sequential acquisition of individual planes, necessitating mechanical or optical adjustments along the z-axis, which inherently limits speed and can introduce artifacts due to motion or misalignment. TranSIM’s innovative beam multiplexing method circumvents these challenges by simultaneously illuminating and imaging multiple planes, drastically reducing the time required for volumetric data acquisition.

### Overall.System.Design.and.Point.Spread.Function.(PSF).Analysis¿

TranSIM incorporates a novel optical architecture designed to achieve simultaneous multi-plane imaging with high spatial and temporal resolution. The schematic in Figure 2 illustrates the key components of the system, including the 488 nm laser source, beam multiplexing optics, and detection paths with multiple imaging sensors for parallel data acquisition. This configuration enables efficient illumination and detection of imaging planes that are both laterally and axially separated. The system leverages precise optical alignment and calibration to maintain uniform intensity and resolution across all imaging planes, ensuring accurate volumetric data capture without axial scanning.

**Figure 1:**
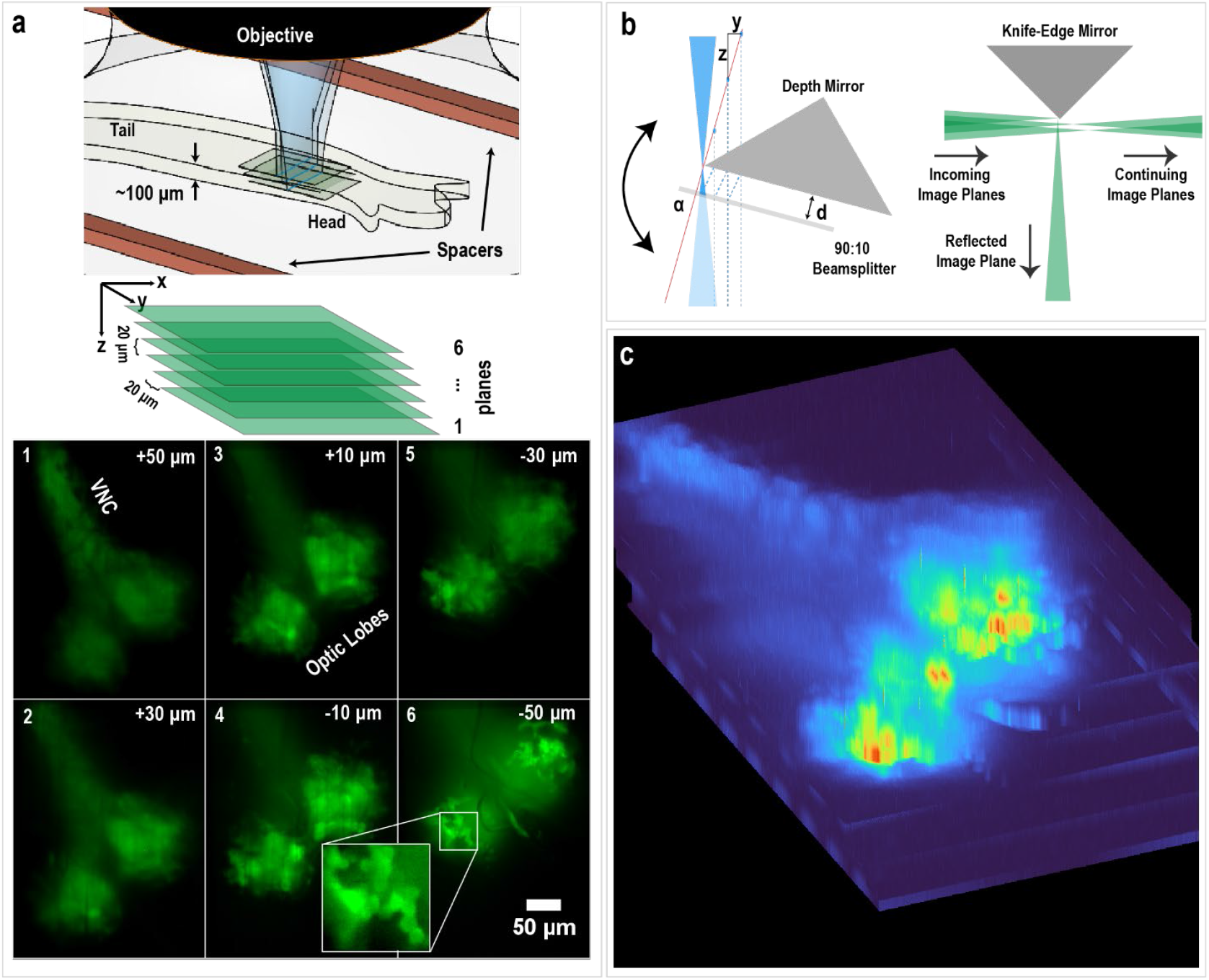
The working principle of Transverse Sheet Illustration Microscopy. A) Six axially and laterally (only three are shown for simplicity) separated beams illuminate the sample creating a parallelepiped volume. Using two sensors, TranSIM captures these six simultaneously imaged planes separated by approximately 20 µm. In the tiled plane display, we see that the first imaging sensor captures the odd pairs of planes (one, three, five), and the second sensor captures the even pairs of planes (two, four, six). Zoom box shows a detailed view of many neurons in the optic lobe. B) To create the illumination beams, a single focused laser beam is spatially multiplexed with a 90:10 beamsplitter paired with a fully reflective mirror cavity. This results in a set of axially and laterally separated beams with separations governed by the angle of the mirror pair and the distance between them. The illumination beams create conjugate fluorescent signals which retain their lateral and axial separation and can therefore be separated using a knife-edge mirror and diverted towards an imaging sensor. C) Three-dimensional reconstruction of the imaging planes after alignment showcasing a Drosophila melanogaster instar 2 larvae brain.

**Figure 2:**
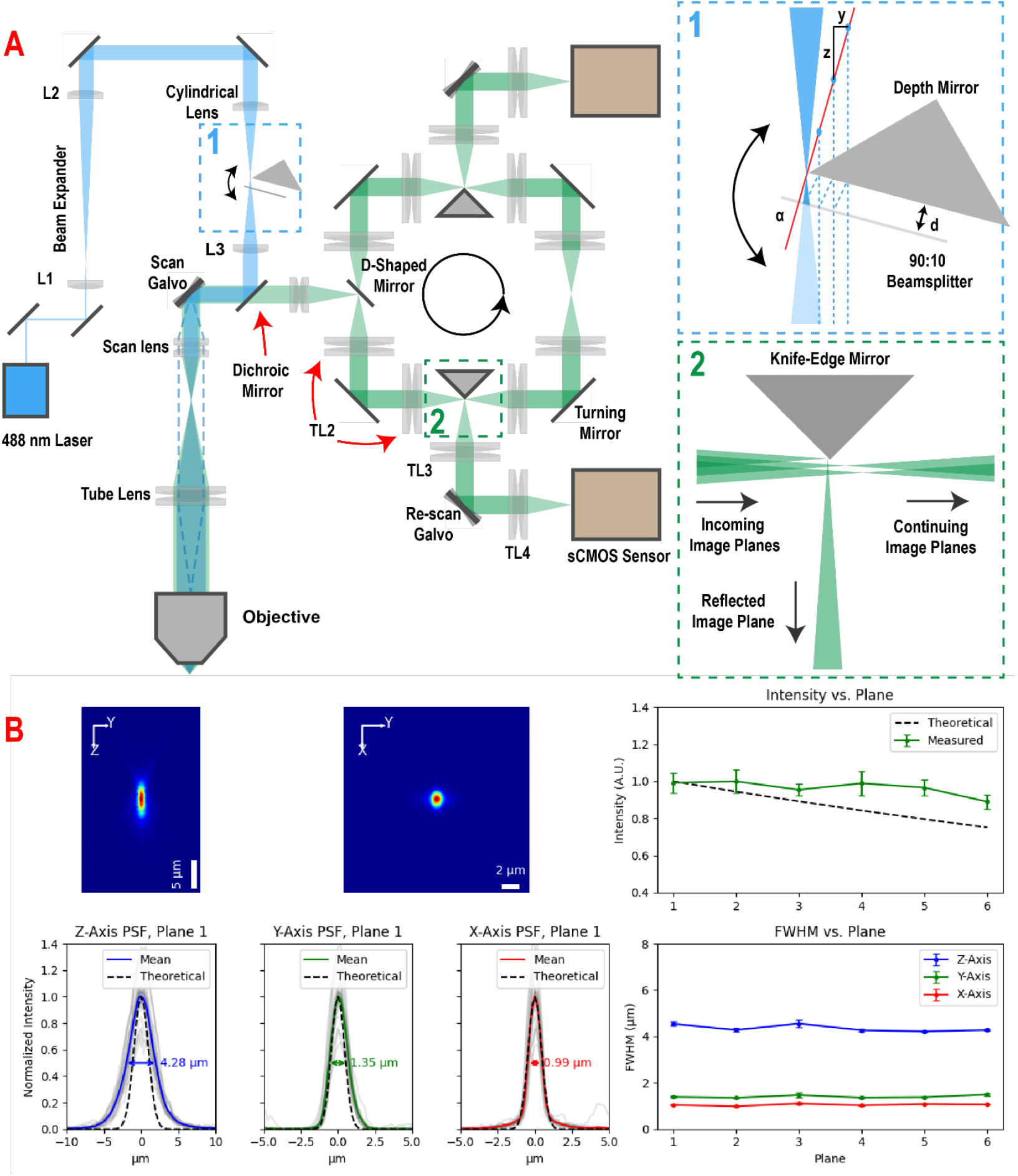
Full schematic of TranSIM and PSF Analysis. A) A 488 nm laser source is beam expanded with a 10X telescope, from a 1/e^2^ beam diameter to a 10/e^2^ beam diameter. The beam is then passed through a cylindrical lens to create the primary scan laser line and multiplexed with a 90:10 beamsplitter and depth mirror. The beams are then diverted to a scanning galvanometer and scanned through a traditional scan-arm configuration. The fluorescent signal returns through the system and de-scanned by the galvanometer and passes through the dichroic mirror. The signal and then relayed through the cycle by four times 4f configuration lens pairs and each of the planes are picked off. The resultant number of planes is three per camera. B) Point spread function and intensity analysis of the system with cross-sectional views and independent XYZ analysis by plane. Utilized 1 µm beads and theoretical PSF was calculated using the modified FWHM taking the bead’s size into account denoted in detail in the methods section.

**Figure 3:**
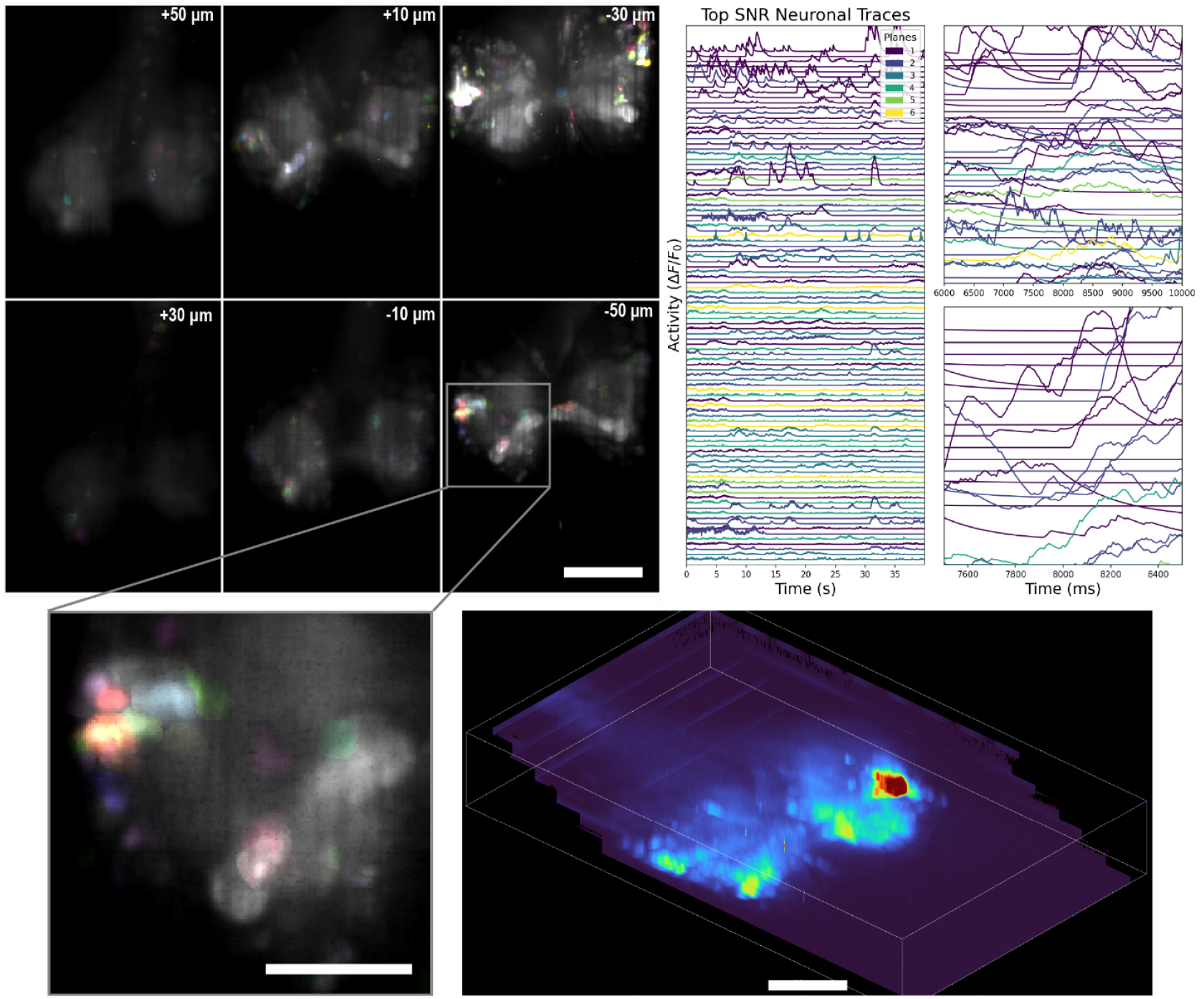
Neural dynamics in Drosophila melanogaster larvae. An instar 2 larva was imaged at 100 volumes per second across 6 planes with an average interplanar distance of 20 µm. The top panel (+30 µm) captures imaging approximately halfway through the optic lobe, while the last plane (−30 µm) was positioned near the ventral side at the bottom of the optic lobe. At the center of the optic lobe, we observed a high density of active neurons, which decreased further into the sample, leaving only a few neurons with a distinct commissure throughout several planes. Scale bar in the tile panel view is 100 µm. Zoomed in to panel +30 µm, we can see high neuron density clustering and not all neurons being selected. Scale bar in zoomed view is 50 µm. The top 100 neuronal traces were plotted over a 40-second time window, with zoomed-in subplots of 8 and 2 seconds to highlight fast neural dynamics. A three-dimensional rendering of the reconstructed volume, visualized via Napari, showcases the alignment of planes and highlights the staircase configuration characteristic of TranSIM. Scale bar in the three-dimensional reconstruction is 50 µm.

To validate the optical performance of TranSIM, we performed a detailed PSF analysis. Figure 2 presents cross-sectional views of the PSF in the XY and YZ planes, demonstrating the system’s resolution in both lateral and axial dimensions. The PSF was measured at multiple imaging planes to evaluate the uniformity of resolution and optical quality across the volume. Gaussian fitting was applied to the data to extract key metrics, including Full-Width-Half-Maximum (FWHM) values. These measurements confirm that TranSIM achieves consistent resolution across all planes, with near-diffraction-limited performance in both the axial and lateral dimensions. The system achieves FWHM values of 4.28 µm in the Z dimension, 1.35 µm in the Y dimension, and 0.99 µm in the X dimension for the first imaging plane and maintains similar resolutions throughout the remaining planes, emphasizing its capability for high-resolution simultaneous volumetric imaging.

In addition to resolution, we quantified the relative intensity drop as a function of plane depth to assess light propagation efficiency through the system. The plot in Figure 2 highlights the variation in intensity across planes, providing insight into the system’s performance in maintaining signal strength. These analyses demonstrate that TranSIM not only provides high-resolution imaging across multiple depths but also ensures uniform signal intensity, which is critical for accurately reconstructing 3D volumes. Together, the schematic and PSF validation data establish the system’s robust optical performance and readiness for high-speed, high-fidelity volumetric imaging.

### Validation.in.Animal.Models¿

To validate TranSIM highspeed three-dimensional imaging capabilities, we imaged Drosophila instar 2 larvae at 100 Hz. We identified more than 200 neurons with significant neuronal activity, which were selected based signal-to-noise (SNR) thresholding rather than morphology alone. The larvae were immobilized with hydrogel and mechanical pressure as described in the methods section. This result shows the system’s ability to maintain both spatial and temporal fidelity during high-speed imaging at 100 Hz. The detection of such many active neurons highlights TranSIM’s robust sensitivity, even in a dynamic, live organism. The simultaneous imaging of multiple planes ensured that neuronal activity across varying depths could be effectively captured without compromising resolution or intensity.

To extract neuronal traces, use utilized the CaImAn Python processing pipeline^43^. Here the tool was able to extract greater than 5000 potential neurons, however, most were rejected based on SNR, enabling us to focus exclusively on well-defined signals. This allowed us to emphasize the temporal properties of TranSIM and to characterize its performance. This analysis revealed clear temporal patterns of neuronal firing, further validating TranSIM’s ability to resolve dynamic neural processes with high precision at significant depths. These results not only highlight TranSIM’s technical advancements but also underscore its potential as a transformative tool for neuroscience research, enabling the exploration of spatiotemporal dynamics and neural connectivity in unprecedented detail.

### Spatiotemporal.Compression¿

Due to TranSIM’s innovative method of de-scanning the image planes, it is possible to manipulate the scanning waveforms to achieve spatiotemporal compression, thereby enhancing the system’s versatility in volumetric imaging. This is accomplished by decoupling the scanning galvanometer’s input amplitude from the rescanning waveform, allowing for effective vertical magnification changes or data compression. Specifically, when a 374 µm amplitude is supplied to the scanning galvanometer, the rescanning waveform can be adjusted to spatially compress the data. This leads to a reduction in the vertical sampling at the sensor, effectively increasing the imaged field of view while maintaining resolution in the x-axis through horizontal binning. This phenomenon, termed spatial compression, provides the ability to capture larger volumetric fields without altering the physical scanning conditions at the sample.

Additionally, TranSIM supports temporal compression, where the rolling shutter region of interest (ROI) at the sensor is reduced, thereby decreasing the number of vertical pixels utilized per frame. By increasing the scanning frequency and reducing the ROI from 920 to 460 pixels, the frame rate can be doubled from 100 Hz to 200 Hz, enabling faster volumetric imaging while preserving the same spatial field of view. This approach not only increases temporal resolution but also reduces the data volume per unit of time, facilitating more efficient data acquisition and processing.

In Figure 4, we illustrate these two types of spatiotemporal compression methods and their impact on volumetric imaging. The top row demonstrates spatial compression, where the vertical scanning amplitude is increased, resulting in an effective vertical binning at the sensor. Using neuronal tracing with the CaImAn processing pipeline, we confirmed that spatial compression enables the capture of fields of view twice as large, increasing the total imaging volume from 374 × 278 × 100 µm^3^ to 748 × 278 × 100 µm^3^. This approach provides a straightforward method for imaging larger biological samples without sacrificing temporal resolution. The bottom row highlights temporal compression, showing that reducing the vertical ROI by half leads to a doubling of the frame rate from 100 Hz to 200 Hz. This capability is particularly advantageous for capturing rapid biological dynamics, such as fast neuronal firing patterns, while maintaining the same volumetric field of view.

**Figure 4:**
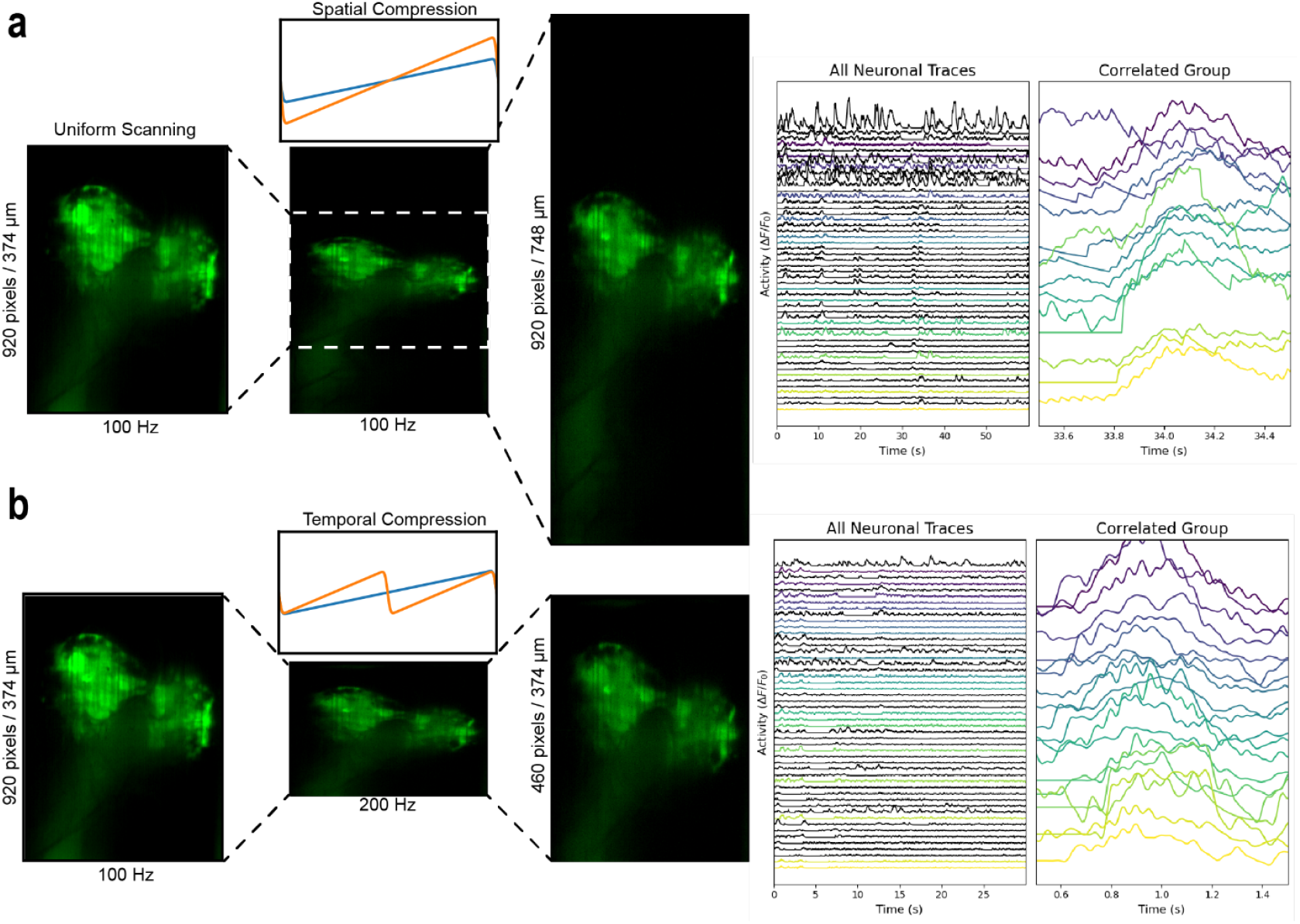
Spatiotemporal compression. Top row, typical 100 Hz volumetric scan can spatially be compressed by increasing the scanning waveform amplitude at the sample. This results in an effective vertical binning at the sensor. The aspect ratio can be regained through horizontal binning. Spatial compression is shown through neuronal tracing with CaImAn to be an effective way of capturing fields of view twice as large, increase the total volume from 374 × 278 × 100 µm^3^ to 748 × 278 × 100 µm^3^. Bottom row, similar shows a 100 Hz volumetric scan being temporally compressed by decreasing the rolling shutter ROI at the sensor from 920 to 460 pixels. This reduction in vertical pixels increases the frame rate from 100 to 200 Hz leading to a temporal data compression while maintaining the same volumetric scan field of view.

Together, these spatiotemporal compression strategies significantly expand the operational flexibility of TranSIM, making it adaptable to a wide range of imaging scenarios. Spatial compression is ideal for applications requiring large fields of view, such as whole-organism imaging or wide-area neuronal activity mapping. Conversely, temporal compression is well-suited for studying high-speed dynamic processes, such as calcium signaling or neural activity, in smaller volumes. By integrating these capabilities, TranSIM achieves a balance of spatial and temporal resolution, addressing key limitations of traditional microscopy systems.

## Discussions

The results presented here demonstrate that TranSIM represents a significant advancement in volumetric imaging, addressing longstanding limitations in traditional microscopy systems. Transverse Sheet Illumination Microscopy provides a framework for rapid and parallel data acquisition. Typically, the fluorescent signal in microscopy is captured with a single sensor, thereby reducing the total bandwidth of the system. Here we have demonstrated that it is possible to spatially parallelize the acquisition of data on the detection side while simultaneously creating a simple illumination schema that can achieve volumetric imaging. Furthermore, we have shown that TranSIM operates at or near diffraction limited resolutions, indicating there to be no significant loss of spatial information as is typical with fast volumetric scanners. TranSIM offers a unique combination of real-time 3D imaging and uniform optical quality across an extended field of view.

We also explored the possibility of utilizing multiple sensors to increase the total bandwidth. Moreover, we configured the system to spatially separate the focal planes the system viewed and translated them to be in focus on these multiple sensors. In addition, by utilizing Hamamatsu Flash 4.0 V2 sensors with a maximum rolling shutter speed limited to 100 kHz with 2048-pixel width, we still reserve the possibility of further improvement by using newer generation sensors such as the Hamamatsu ORCA-Quest 2 qCMOS versions with overall greater bandwidths. For example, the qCMOS has a 120 kHz line rate with 4096 pixels, which would provide an overall increase of 2.4x bandwidth, increasing the 0.38 GHz pixel rate 0.912 GHz. Additionally, TranSIM’s design does not explicitly call for the use of two-dimensional sensors for imaging. Since the system inherently de-scans the image planes before entering the realignment cycle, theoretically, we can use line sensors. This approach could result in reduced total costs and system complexity and is something that we aim to explore in the future.

Furthermore, TranSIM’s design can be coupled with other systems mentioned in this work, particularly those of the 2-photon variety, like Light Bead Microscopy and Reverberation 2P Microscopy, as they are already tuned for temporal multiplexing. By combining TranSIM’s detection side multi-sensor approach, we can overcome the overall bandwidth that does systems have. We expect TranSIM to pave the way towards the parallel data acquisition on the detection side. By providing rapid and detailed volumetric data, TranSIM opens new avenues for exploring neural connectivity and spatiotemporal dynamics in ways that were previously unattainable.

## Materials and Methods

To achieve simultaneous imaging of multiple axially separated planes, TranSIM generates laterally and axially separated illumination beams on the illumination side. These beams are scanned laterally using a galvanometer and relayed through the system to the imaging plane in front of the objective. On the detection side, the samples fluoresce, the light is captured using the same objective, and it propagates through the same scanning relay to the galvanometer in reverse relative to the illumination. The planes are subsequently de-scanned by the galvanometer. Once de-scanned, the imaging planes are split from the illumination using a dichroic mirror which allows the planes into an optical resonator. This optical resonator focuses and realigns the imaging planes on the multiple imaging sensors. Here we detail the methods we employ regarding the various parts of TranSIM, the illumination, detection, system control, and custom software. We detail how we perform point spread function (PSF) analysis using fluorescent beads and biological specimen data analysis. Lastly, we covered the sample preparation of the biological systems, namely Drosophila. melanogaster?we tested TranSIM.

### Illumination.System.Design¿

TranSIM is a 1P system using a 488 nm, 100 mW Coherent Sapphire (Coherent Inc.) continuous wavelength laser. The laser beam diameter is expanded using a −50 mm (LC1715-A, Thorlabs) to 250 mm (LA1461-A, Thorlabs) telescope for a 5 times beam expansion from a 1 mm to 5 mm diameter. The beam is then passed through a 100 mm cylindrical lens (LJ1567RM, Thorlabs) to create the prime laser line that will be multiplexed. For multiplexing, a 90:10 R:T beamsplitter (BS) (BSX10R, Thorlabs) is placed immediately after the focal line of the cylindrical lens with angle α from the optical axis. A wedge mirror is then brought it a close as possible to the laser line without clipping it and placed parallel to the BS. The BS allows 10% of the light through and 90% is reflected to the wedge mirror which in turn fully reflects the beam back towards to the BS. Here the beam again passes through the BS with 10% of the remaining power and the process repeats. With each subsequent bounce, the transmitted beam’s power decreases exponentially (*Pn*= 0.1 ×(0.9)^*n*−1^×*P*_0_). Nevertheless, this results in an infinite set of beams that are equally laterally and axially separated. The lateral separation is introduced by the slight angle α, and the axial separation is introduced by the spatial delay introduced by the bouncing to the next lens.

The multiplexed beams are then collected with a 100 mm achromatic lens (AC254-100-A, Thorlabs) off centered such that the Nth/2 plane (of used planes on the detection side) is placed at the centered axis. The beams are then reflected off a Dichroic mirror (T495lpxru, Chroma) towards a galvanometer for scanning. The scan lens is an effective 100 mm focal length telecentric lens created using a pair of 200 mm achromatic lenses (ACT508-200-A, Thorlabs). Similarly, the tube lens is a 200 mm effective focal length lens comprised of a pair of 400 mm achromatic lenses (ACT508-400-A, Thorlabs). Telecentric lenses were chosen to maintain constant magnification. For our imaging lens we chose a Nikon 16X 0.8 NA objective. The laser lines are then scanned laterally using the galvanometer.

### Detection.System.Design¿

The detection beams are collected with the same Nikon 16X 0.8 NA objective and are initially scanned in the same manner as the illumination beams until they are de-scanned by the galvanometer. Once de-scanned, the beams are then passed through the dichroic mirror where they are injected into the cyclic module. The initial insertion is made by focusing the focal lines using a 100 mm effective focal length telecentric lens using a pair of 200 mm achromatic lenses (AC254-200-A, Thorlabs) onto a D-shaped pickoff mirror (BBD05-E02, Thorlabs). The beams are initially laterally offset in the cyclic module as this will allow space for realignment and focusing.

Now inside of the cyclic module, the beams are relayed using a 4f relays composed of a 100 mm effective focal length telecentric lenses made from pairs of 200 mm lenses (ACT508-200-A, Thorlabs) and a mirror for 90-degree reflection placed between the telecentric lenses. At the relayed focal plane, a knife-edge mirror (MRAK25-P01, Thorlabs) is brought in close enough to the beams to pick-off the first image plane. This image plane is then relayed and rescanned into an ORCA-Flash4.0 V2 sCMOS sensor (Hamamatsu) with the rolling shutter enabled. The rescanning is produced by a galvanometer. The remaining beams are relayed with a pair of identical 4f telecentric relays to the second knife-edge pick-off mirror where the second beam is sent to the sCMOS sensor. Using the last 4f relay, the remaining beams are laterally adjusted by physically translating the 4f relay to be offset from the previous 3 relays. The focusing is achieved by adjusting the position of the 4f relay lens distances. The beams are then allowed to pass underneath the D-shaped pick-off mirror with the beams having been realigned and refocused to be adjacent to the initial pass and the cycle repeats.

### Systems.Control¿

To synchronize the illumination and detection components of TranSIM, we developed a custom Python software suite. The suite’s first module generates waveforms to synchronize the electro-optical-mechanical devices at the hardware level. The system consists of two sCMOS sensors, one scanning galvanometer, one re-scanning galvanometers (one per camera), and a 10 nm precision piezo actuator (P-721, Physik Instrumente). The data acquisition (DAQ) workstation controls these devices using the analog output of an NI PCIe 6363 (Texas Instruments) DAQ card operating with a sampling rate (*f*_*s*_) of 200 kSamples/second. The high sampling rate is chosen to avoid waveform aliasing. The sCMOS sensors are externally triggered using a 50% duty cycle square waveform with frequency (*f*_*w*_) depending on the ROI used. For example, we commonly operate the system with frequencies of 10, 50, 100, and 200 Hz. With the same frequency as the camera, the scanning galvanometer is supplied with a low pass filtered triangle waveform. The waveform is synthesized based on specified parameters (frequency, amplitude, phase, offset, duty cycle) and smoothed using a Gaussian filter and a 98% duty cycle. This approach effectively attenuates high-frequency noise, yielding a controlled output signal with defined frequency characteristics. For the specified waveforms, we utilized a cut-off frequency (*f*_*c*_) approximately double the scanning frequency, *f*_*c* ≈_ 2*f*_*w*_. The rescanning galvanometers undergo a similar waveform parametrization procedure. However, due to variability in the manufacturing of these devices, the line-scanning speed produced by the galvanometers is synchronized with the respective camera by adjusting the internal line interval speed within the camera parameters. Communication with the laser is established via a serial connection, allowing precise control over laser operations directly through the application.

### Data.Acquisition¿

The second module in the Python suite controls the data acquisition of the system. The ORCA-Flash4.0 V2 sCMOS sensors are connected to the DAQ via Camera Link PCIe cards (FBD-1XCLD-2PE8, Active Silicon). The data is then processed by the DAQ using custom Python software that allows for real-time visualization of the data via multi-threaded processing and OpenCV display. The software also allows simultaneous display and recording of the data and is fully synchronized with the systems control software. Data is transferred to and stored directly to NVMe drives in a raw and uncompressed h5 file format.

### Point.Spread.Function.Analysis¿

For system characterization, 1 µm microspheres (TetraSpeck, ThermoFisher) were imaged. The beads were ultrasonicated for 10 minutes to remove dense particulates. After ultrasonication, 5 µL of 2.0 × 10^10 particles/mL concentration were added to 50 µL of 2% agarose W/V solution and mixed thoroughly. The new concentration of 1.82 × 10^9 particles/mL was then transferred and placed on a glass slide with 150 µm spacers. A cover glass was added to create a flat imaging surface.

To characterize the axial resolution of the TranSIM system, we imaged a calibration sample at a 100 Hz framerate while performing an axial scan using a piezo actuator with a 0.5 µm step size. This fine step size was chosen to adequately sample the system’s point spread function (PSF) along the z-axis, ensuring high precision in axial resolution measurements. A total of 150 frames were captured, covering a 75 µm axial range, which remains well within the piezo actuator’s operational limits, minimizing hysteresis effects. The acquired data was processed using custom Python scripts to extract critical parameters describing the system’s optical performance. Below, we outline the detailed pipeline for processing and analyzing the data.

The first step in the pipeline involves loading the raw data captured by both cameras into memory and splitting the three-dimensional data stack into a four-dimensional hyperstack. This hyperstack is generated by subdividing each raw camera frame into N planes, corresponding to the system’s multi-plane imaging configuration. Plane-to-plane image registration was then performed using the pyStackReg library, which aligns the datasets and removes any lateral drift introduced during imaging. This step ensures consistency across all planes, preserving the spatial coherence necessary for accurate PSF analysis. Subsequently, we identified the locations of fluorescent beads by applying a binarization algorithm to detect the brightest intensity points in the image stacks. Cross-sectional slices of these beads were extracted in all three dimensions (x, y, z), centering the slices on the bead’s brightest point for further analysis.

The PSF of the system was characterized by fitting a three-dimensional Gaussian function to the intensity profile of each bead. The general form of the Gaussian profile is:

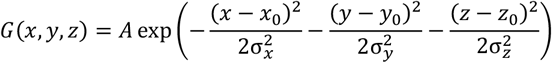

where *G*(*x, y, z*) represents the intensity distribution, A is the peak amplitude, (*x*_0_, *y*_0_, *z*_0_) are the coordinates of the center of the bead, and *σ*_*x*_, *σ*_*y*_, *σ*_*z*_ are the standard deviations of the Gaussian distribution along the *x, y, and z* axes, respectively. From this profile, the FWHM for each dimension is calculated using the relationship:

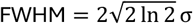

These calculations provide a quantitative measure of the system’s resolution in *laterall* (*x, y*) *axial* (*z*) dimensions. The theoretical resolution limits are defined by the following equations, which depend on the wavelength (*λ*) of the light source and the numerical aperture (*NA*) of the objective lens:

- *Laterall* (*x, y*) resolution

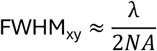
- *Axial* (*z*) resolution

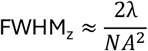

The theoretical resolution of the TranSIM system was determined by considering the detection numerical aperture (NA) and the size of the fluorescent beads used as calibration targets. The lateral and axial FWHM values were adjusted to account for the additional convolution introduced by the finite size of the beads. These corrections ensure that the calculated resolution reflects the system’s performance more accurately.

For lateral resolution, the detection NA and detection wavelength were used to compute the theoretical lateral FWHM. The bead size (1.0 µm) was included in the calculation as an additive term to account for its contribution to the observed intensity profile. The lateral FWHM was calculated as:

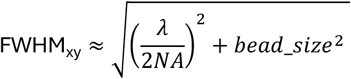

For axial resolution, the system’s detection NA and detection wavelength were used alongside the bead size. The axial FWHM was computed as:

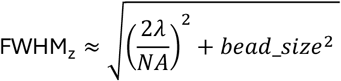

These modified equations provide a more realistic estimation of the system’s resolution by integrating the bead size and the effects of detection NA. This approach ensures that the observed PSF measurements accurately reflect the underlying optical performance of the system.

To evaluate the PSF across the entire imaging volume, we performed this analysis independently for each plane in the hyperstack, yielding a multi-dimensional PSF dataset: *PSF*(*x, y, z, plane*). Additionally, the relative intensity across planes was analyzed to quantify signal attenuation as a function of depth. This comprehensive characterization provides key insights into the system’s optical performance, verifying its ability to maintain high resolution and uniform intensity across all imaging planes.

### Inter=Planar.Distance.and.Volumetric.Field=of=View¿

TranSIM produces data with a stair-case profile, leading to a parallelepiped-like volume. The lateral and axial offsets are determined by the angle of incidence, *α*, on the 90:10 beamsplitter, and the separation distance to the depth mirror, *d*. Pre-magnification lateral plane separation, *y*, is determined by the following trigonometric relation:

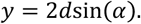

Similarly, the pre-magnification axial displacement, *z*, determined by the following trigonometric relation:

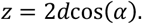

After effective system magnification, *M*_*eff*_, the lateral and axial distances change inversely linearly and quadratically, respectively^44^. The modified magnified lateral and axial plane separations are:

- *Lateral* (*y*).plane separation

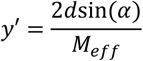
- *Axial* (*z*).plane separation

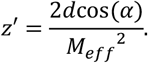

Here, we can define the magnified angle, *α*′, determining the parallelepiped shearing:

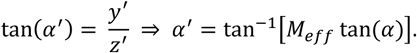

Now we calculate the volumetric FOV using the parallelepiped volume equation:

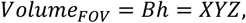

And

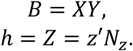

Where *B*, is the base area of an image plane, also defined as *XY* and *h* is the total volume height, also defined as *Z*, where the total height is the number of planes, *N*_*z*_ times the magnified axial plane separation, *z*′. Because of the inversely quadratic dependence on magnification for the magnified angle *α*′, we see that to create a usable parallelepiped volume, namely one with an angle *α*′ ≤ 45^°^, we must restrict our angle of incidence to *α* ≤ 3.58^°^.

### Drosophila.melanogaster.Larvae.Sample¿

Fruit flies were maintained at 25°C with 40% humidity under a 12-hour light/dark cycle, following standard husbandry practices^45^. To ensure high indicator protein expression, new parent flies were selected every 10–14 days from the youngest bottles containing recently emerged adults. Flies were anesthetized in a 4°C refrigerator for one hour, sex-differentiated under a microscope on a cold plate, and then placed into media containers (20 females and 10 males per bottle; 18 females and 9 males per vial). Parent flies were transferred to fresh media bottles as needed for continuous maintenance.

To monitor calcium-dependent neural activity in larvae, we used genetically encoded calcium indicators (GECIs), specifically GCaMP6f, expressed under the GAL4/UAS system^46^. Panneuronal expression was achieved by crossing female flies carrying the UAS-GCaMP6f construct with males of the R57C10-GAL4 driver line (also referred to as nSyb-GAL4)^47^. GCaMP6f was chosen for its high signal-to-noise ratio, fast kinetics, and sensitivity, enabling the detection of rapid calcium transients in neurons.

For immobilization, we employed a combination of mechanical, cold anesthesia, and chemical methods to optimize stability during imaging. First, a 25% (w/v) Pluronic F127 (PF127) hydrogel solution was prepared to serve as a mechanical restraint at room temperature^48^. Second, HL3.1 saline solution, a widely used medium for Drosophila larvae, was prepared as a rinsing and buffer medium^49^. Finally, larvae were anesthetized using controlled exposure to diethyl ether fumes, based on the method described by Kakanj et al^50^.

The immobilization process began by selecting second-instar larvae (4–5 days after egg laying). Larvae were extracted from the media and placed on a glass slide equipped with 150-µm copper tape spacers. The slide was then placed in a chamber containing approximately 10 mL of diethyl ether at a distance of ∼7 cm and stored at 4°C, combining chemical and cold anesthesia. After 30 minutes, the slide was removed, and each larva was rinsed with 0.5 µL of HL3.1 solution to remove residual media. Next, 20 µL of PF127 hydrogel was applied, followed by the gentle placement of an 18 × 18 mm coverslip (Corning, 41121800) to provide mechanical pressure while enhancing brain visibility. The 150-µm spacers ensured uniform compression, preventing excessive force that could damage the specimen. The prepared slides were then placed on a 25°C heating plate for 2–3 minutes to accelerate hydrogel solidification. Following this procedure, larvae were fully immobilized and ready for calcium imaging.

## Acknowledgements

This work was supported with funding from NIH grant R21EY028395. We also acknowledge the use of reagents at the Advanced Light Microscopy/Spectroscopy Laboratory (RRID: SCR_022789) and the Leica Microsystems Center of Excellence at the California NanoSystems Institute at UCLA. Additionally, we extend our appreciation to the Elegant Mind Club at UCLA, with special thanks to David Reynolds, Eliana Artenyan, Sandra Karon, and Altamash Mahsud for their invaluable contributions to animal husbandry and sample preparation.

## References

1. Miyawaki, A. et.al¡ Fluorescent indicators for Ca2+based on green fluorescent proteins and calmodulin. Nature 388, 882–887 (1997).

2. Chen, T.-W. et.al¡ Ultrasensitive fluorescent proteins for imaging neuronal activity. Nature 499, 295–300 (2013).

3. Nakai, J., Ohkura, M. & Imoto, K. A high signal-to-noise Ca2+ probe composed of a single green fluorescent protein. Nat.Biotechnol 19, 137–141 (2001).

4. Dana, H. et.al¡ High-performance calcium sensors for imaging activity in neuronal populations and microcompartments. Nat.Methods 16, 649–657 (2019).

5. Daria, V. Spatio-temporal parameters for optical probing of neuronal activity. (2021).

6. Kerr, R. et.al¡ Optical Imaging of Calcium Transients in Neurons and Pharyngeal Muscle of C¡. elegans. Neuron 26, 583–594 (2000).

7. Tian, L. Imaging neural activity in worms, flies and mice with improved GCaMP calcium indicators. (2009).

8. Dana, H. et.al¡ Sensitive red protein calcium indicators for imaging neural activity. eLife 5, e12727 (2016).

9. Petreanu, L. Activity in motor–sensory projections reveals distributed coding in somatosensation. (2012).

10. Ji, N., Freeman, J. & Smith, S. L. Technologies for imaging neural activity in large volumes. Nat. Neurosci 19, 1154–1164 (2016).

11. Mertz, J. Strategies for volumetric imaging with a fluorescence microscope. Optica?.OPTICA 6, 1261–1268 (2019).

12. Peng, H. et.al¡ BrainAligner: 3D Registration Atlases of Drosophila Brains. Nat.Methods 8, 493–500 (2011).

13. Scheffer, L. K. et.al¡ A connectome and analysis of the adult Drosophila central brain. eLife 9, e57443 (2020).

14. Dorkenwald, S. et.al¡ Neuronal wiring diagram of an adult brain. Nature 634, 124–138 (2024).

15. Zheng, Z. et.al¡ A Complete Electron Microscopy Volume of the Brain of Adult Drosophila. melanogaster. Cell 174, 730-743.e22 (2018).

16. Dunsby, C. Optically sectioned imaging by oblique plane microscopy. Opt¡.Express?.OE 16, 20306–20316 (2008).

17. Zhang, L., Capilla, A., Song, W., Mostoslavsky, G. & Yi, J. Oblique scanning laser microscopy for simultaneously volumetric structural and molecular imaging using only one raster scan. Sci. Rep 7, 8591 (2017).

18. Kim, J. Recent advances in oblique plane microscopy. Nanophotonics 12, 2317–2334 (2023).

19. Voleti, V. et.al¡ Real-time volumetric microscopy of in vivo dynamics and large-scale samples with SCAPE 2.0. Nat.Methods 16, 1054–1062 (2019).

20. Denk, W., Strickler, J. H. & Webb, W. W. Two-Photon Laser Scanning Fluorescence Microscopy. Science 248, 73–76 (1990).

21. So, P. T. C., Dong, C. Y., Masters, B. R. & Berland, K. M. Two-Photon Excitation Fluorescence Microscopy. Annual.Review.of.Biomedical.Engineering 2, 399–429 (2000).

22. Amir, W. et.al¡ Simultaneous imaging of multiple focal planes using a two-photon scanning microscope. Opt¡.Lett¡?.OL 32, 1731–1733 (2007).

23. Cheng, A., Gonçalves, J. T., Golshani, P., Arisaka, K. & Portera-Cailliau, C. Simultaneous two-photon calcium imaging at different depths with spatiotemporal multiplexing. Nat.Methods 8, 139–142 (2011).

24. Beaulieu, D. R., Davison, I. G., Kılıç, K., Bifano, T. G. & Mertz, J. Simultaneous multiplane imaging with reverberation two-photon microscopy. Nat.Methods 17, 283–286 (2020).

25. Demas, J. et.al¡ High-speed, cortex-wide volumetric recording of neuroactivity at cellular resolution using light beads microscopy. Nat.Methods 18, 1103–1111 (2021).

26. Akerboom, J. et.al¡ Optimization of a GCaMP Calcium Indicator for Neural Activity Imaging. J¡. Neurosci¡ 32, 13819–13840 (2012).

27. Levoy, M., Ng, R., Adams, A., Footer, M. & Horowitz, M. Light field microscopy. ACM.Trans¡. Graph¡ 25, 924–934 (2006).

28. Levoy, M., Zhang, Z. & Mcdowall, I. Recording and controlling the 4D light field in a microscope using microlens arrays. Journal.of.Microscopy 235, 144–162 (2009).

29. Prevedel, R. et.al¡ Simultaneous whole-animal 3D-imaging of neuronal activity using light-field microscopy. Nat.Methods 11, 727–730 (2014).

30. Levoy, M. & Hanrahan, P. Light field rendering. in Proceedings.of.the.89rd.annual.conference.on. Computer.graphics.and.interactive.techniques 31–42 (Association for Computing Machinery, New York, NY, USA, 1996). doi:10.1145/237170.237199.

31. Pégard, N. C. et.al¡ Compressive light-field microscopy for 3D neural activity recording. Optica?. OPTICA 3, 517–524 (2016).

32. Lu, Z. et.al¡ Phase-space deconvolution for light field microscopy. Opt¡.Express?.OE 27, 18131–18145 (2019).

33. Guo, C., Liu, W., Hua, X., Li, H. & Jia, S. Fourier light-field microscopy. Opt¡.Express?.OE 27, 25573–25594 (2019).

34. Bai, L. et.al¡ Volumetric voltage imaging of neuronal populations in the mouse brain by confocal light-field microscopy. Nat.Methods 21, 2160–2170 (2024).

35. Badon, A. et.al¡ Video-rate large-scale imaging with Multi-Z confocal microscopy. Optica?. OPTICA 6, 389–395 (2019).

36. Tsang, J.-M. et.al¡ Fast, multiplane line-scan confocal microscopy using axially distributed slits. Biomed¡.Opt¡.Express?.BOE 12, 1339–1350 (2021).

37. Weber, T. D., Moya, M. V., Kılıç, K., Mertz, J. & Economo, M. N. High-speed multiplane confocal microscopy for voltage imaging in densely labeled neuronal populations. Nat.Neurosci 26, 1642–1650 (2023).

38. Sheppard, C. J. R. & Mao, X. Q. Confocal Microscopes with Slit Apertures. Journal.of.Modern. Optics 35, 1169–1185 (1988).

39. Im, K.-B., Han, S., Park, H., Kim, D. & Kim, B.-M. Simple high-speed confocal line-scanning microscope. Opt¡.Express?.OE 13, 5151–5156 (2005).

40. Wolleschensky, R., Zimmermann, B. & Kempe, M. High-speed confocal fluorescence imaging with a novel line scanning microscope. JBO 11, 064011 (2006).

41. Baumgart, E. & Kubitscheck, U. Scanned light sheet microscopy with confocal slit detection. Opt¡.Express?.OE 20, 21805–21814 (2012).

42. Mei, E., Fomitchov, P. a., Graves, R. & Campion, M. A line scanning confocal fluorescent microscope using a CMOS rolling shutter as an adjustable aperture. Journal.of.Microscopy 247, 269–276 (2012).

43. Giovannucci, A. et.al¡ CaImAn an open source tool for scalable calcium imaging data analysis. eLife 8, e38173 (2019).

44. Born, M. & Wolf, E. Principles.of.Optics¿.Electromagnetic. Theory.of.Propagation?.Interference. and.Diffraction.of.Light. (Cambridge University Press, Cambridge, 1999). doi:10.1017/CBO9781139644181.

45. Mołoń, M. Effects of Temperature on Lifespan of Drosophila melanogaster from Different Genetic Backgrounds: Links between Metabolic Rate and Longevity. (2020).

46. Brand, A. H. Targeted gene expression as a means of altering cell fates and generating dominant phenotypes. (1993).

47. Pfeiffer, B. Tools for neuroanatomy and neurogenetics in Drosophila. (2008).

48. Dong et. al. Reversible and long-term immobilization in a hydrogel-microbead matrix for high-resolution imaging of Caenorhabditis elegans and other small organisms. (2018).

49. Feng et. al. A MODIFIED MINIMAL HEMOLYMPH-LIKE SOLUTION, HL3.1, FOR PHYSIOLOGICAL RECORDINGS AT THE NEUROMUSCULAR JUNCTIONS OF NORMAL AND MUTANT DROSOPHILA LARVAE. (2004).

50. Kakanj, P., Eming, S. A., Partridge, L. & Leptin, M. Long-term in vivo imaging of Drosophila larvae. Nat.Protoc 15, 1158–1187 (2020).

